# Nucleosides rescue replication-mediated genome instability of human pluripotent stem cells

**DOI:** 10.1101/853234

**Authors:** Jason A. Halliwell, Thomas J. R. Frith, Owen Laing, Christopher J Price, Oliver J. Bower, Dylan Stavish, Paul J. Gokhale, Zoe Hewitt, Sherif F. El-Khamisy, Ivana Barbaric, Peter W. Andrews

## Abstract

Human pluripotent stem cells (PSC) often acquire genetic changes on prolonged culture, which pose concerns for their use in research and regenerative medicine (Amps et al., 2011, Seth et al., 2011). The acquisition of these changes during culture necessarily first requires mutation and then selection of those mutations that provide a growth advantage. Whilst selection accounts for the recurrent nature of the variants commonly reported (Draper et al., 2004, Olariu et al., 2010), the mechanisms of mutation in PSC remain largely elusive. Here we show that, in contrast to somatic cells, human PSC have an increased susceptibility to DNA damage and mitotic errors, both of which are caused by heightened replication stress in PSC and this can be alleviated by culture with exogenous nucleosides. These results reflect the requirement for rapid replication of human PSC enabled by a truncated G1 (Becker et al., 2006, Becker et al., 2010) that impairs the preparation of these cells for the ensuing DNA replication. A similar relationship has been shown in relation to chromosomal instability in cancer cells (Burrell et al., 2013, Wilhelm et al., 2019) but PSC differ by replication stress triggering apoptosis (Desmarais et al., 2012, Desmarais et al., 2016). Nevertheless, evasion of this response still leads to the appearance of genetic variants that are of concern for regenerative medicine. The inclusion of nucleosides into culture media greatly improves the efficiency of human PSC culture and minimises the acquisition of genomic damage.

---

DNA double strand breaks (DSBs) are a particularly detrimental type of DNA damage. Unrepaired, or erroneously repaired, DSBs can jeopardize genome stability by leading to mitotic aberrations and structural and chromosomal instability (Ichijima et al., 2010, Janssen et al., 2011), like those frequently observed in human PSC (Amps et al., 2011). The human induced PSC line MIFF1, herein referred to as hiPSC1, exhibited an increased number of gH2AX foci, known to mark sites of DSB, compared to the fibroblast from which it was reprogrammed and to its differentiated derivatives obtained by treatment with CHIR99021 for 5 days (**Fig. 1a and Supplementary Fig. 1**). Two other PSC lines that we examined, TC113 and MShef11, referred to in the figures as hiPSC2 and hESC respectively, showed a similarly elevated level of DNA damage compared to their differentiated derivatives (**Fig. 1a and Supplementary Fig. 1**). These observations were confirmed by a neutral comet assay that showed an increased tail moment in the undifferentiated cells in each case (**Fig. 1b**), demonstrating that a high level of genome damage is associated with the pluripotent state and decreases upon differentiation.

**Figure 1.**
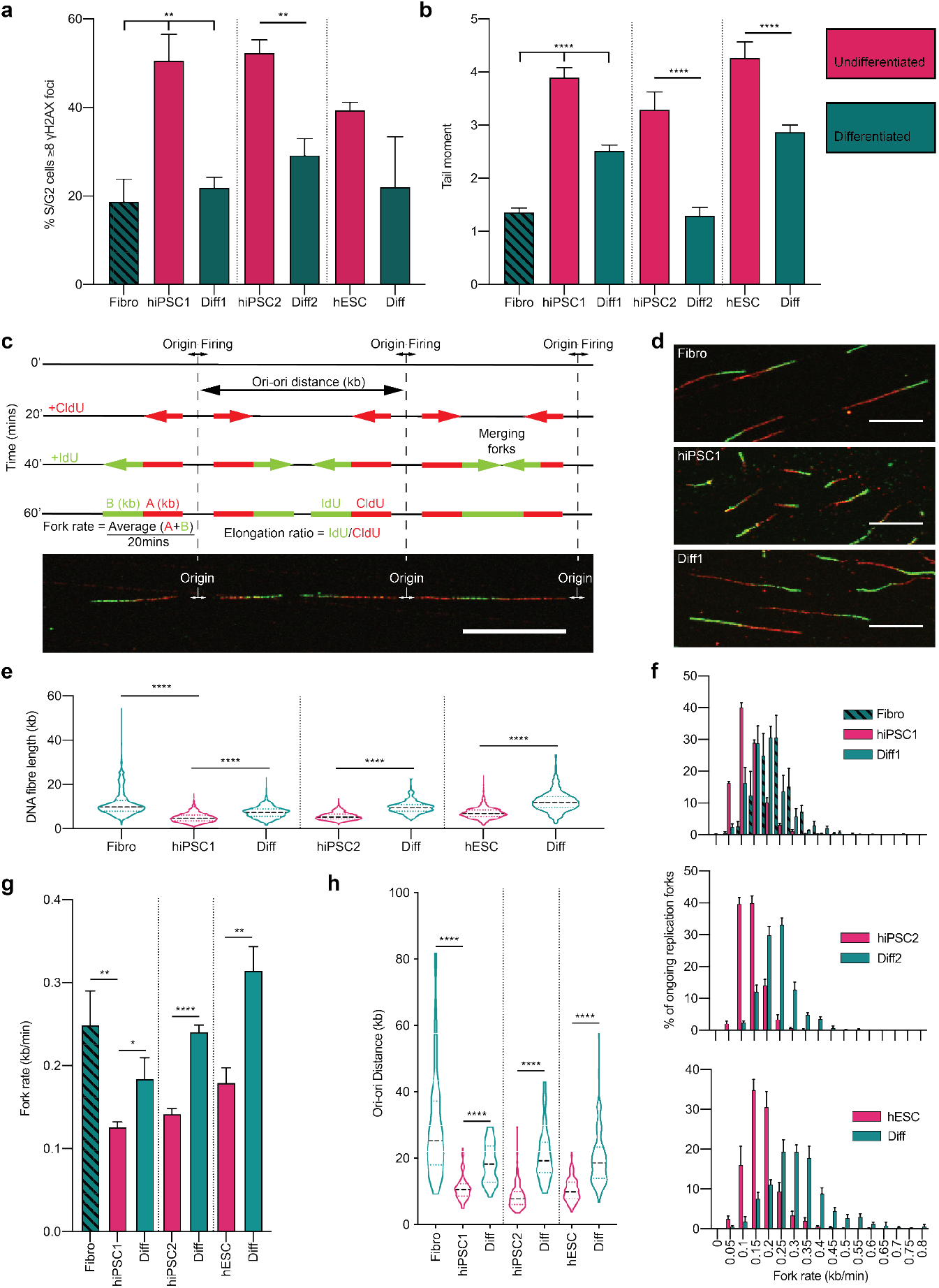
Replication stress and a susceptibility to DNA damage is a characteristic of undifferentiated human PSC. a, b, Double strand breaks observed in human PSCs compared to a parent fibroblast line (Fibro) of MIFF1 (hiPSC1) and following differentiation (Diff) of each human PSC line; MIFF1 (hiPSC1), TC113 (hiPSC2) and mShef11 (hESC). **a**, Frequency of S/G2 cells with ≥8 gH2AX foci. The S/G2 phase was determined from nuclear DNA content. Data in **a** are mean values of 3 independent experiments ± s.d., two-tailed *t*-test, **P<0.01, (*n* > 100 cells per cell line per experiment). t-values; 7.031, 7.781, 8.293 and 2,605 (left to right) **b**, Average tail moment from neutral comet assays. Data displayed is from 3 independent experiments ± s.e.m., two-tailed *t*-test, ****P<0.0001, (*n* ≥ 300 cells per cell line per experiment). t-values; 12.253, 6.484, 5.574 and 4.317. **c-h**, DNA fibre assays were performed; Combined CldU/IdU fibre length, elongation ration CldU:IdU, replication rates and replication origin distances of individual replication forks was assessed. Comparisons were made between three human PSC and their isogenic somatic counterparts. **c**, Schematic of DNA fibre analysis. Sequential 20minute pulses of CldU and IdU labelled the progressing replication forks. Measurements of CldU and IdU lengths facilitate the analysis of replication fork dynamics. **d**, Representative DNA fibres are shown for Fibro, hiPSC1 and Diff. Scale bar, 10μm. **e**, Combined length of CldU and IdU in individual fibres (*n* > 200 forks per cell line per experiment, *n* = 3 experiments). Median distance, 25^th^ and 75^th^ quartiles are presented, two-tailed *t*-test, ****P<0.0001. t-vales; 39.65, 25.28, 36.68 and 30.43 (left to right) **f**, Distribution of replication fork rates (*n* > 200 forks per cell line per experiment, *n* = 3 experiments), data is mean value from each experiment ± s.e.m. **g**, Mean fork rates from (**f**) ± s.d., two-tailed *t*-test, *P<0.05, **P<0.01, ****P<0.0001, (*n* = 3 experiments). t-values; 5.073, 3.780, 15.750 and 6.789 (left to right). **h**, Distribution of adjacent origins distance measurements (Ori-ori). Median distance, 25^th^ and 75^th^ quartiles are presented, two-tailed *t*-test, ****P<0.0001 (*n* > 30 per cell line, *n* = 3 experiments). t-values; 8.772, 7.348, 17.650, 11.380 (left to right).

A common cause of DSB during S phase in cancer cells is the slowing, stalling and collapse of replication forks, recognised as DNA replication stress (Ichijima et al., 2010). To analyse the replication dynamics in undifferentiated and differentiated cells, we utilised the DNA fiber assay. Here, the newly synthesized DNA is pulse labelled successively with thymidine analogues cholorodeoxyuridine (CldU) and iododeoxyuridine (IdU) for 20 minutes each, and then visualized by fluorescently labeled antibodies (**Fig. 1c**). By measuring the total length of the CldU and IdU labelling in each fibre, we found a decrease in the length of newly synthesised fibres in the undifferentiated state (**Fig. 1c-e).** Replication fork speed, calculated by measuring the average length of labelled fibres, was significantly slower in undifferentiated PSC compared to their isogenic somatic counterparts **(Fig. 1c,f,g)**. Further, we found an increase in the abundance of origins of DNA replication as evidenced by a decrease in replication origin-to-origin distance (Ori-ori) in PSC **(Fig. 1c,h)**. Overall, these results show that DNA replication in pluripotent cells is considerably perturbed, predisposing them to DNA damage, notably DSB.

The association of genome damage with the pluripotent rather than the somatic state of the same cell line, suggests that features linked to pluripotency impart replication stress on PSC. A pertinent key difference between the pluripotent and somatic cell state is the rapid progression of PSC through G1, driven by atypical expression of cyclins (Becker et al., 2006), Ahuja et al., 2016). By time-lapse microscopy and EdU pulse chase analysis we found that the human PSC line, hiPSC1, exhibited a reduced cell cycle time when compared to its parent fibroblast line **(Fig. 2a)**. Specifically, the abbreviated cell cycle time was solely due to a truncated G1 **(Fig. 2a)**. Consistent with the reduction in the length of G1, Cyclin D2 (CCND2) and Cyclin E (CCNE1 and CCNE2), which are known to drive the Rb-E2F pathway and allow rapid progression of cells through G1 (Hinds et al., 1992, Lundberg and Weinberg, 1998), were highly expressed in the undifferentiated hiPSC1 compared to the corresponding parent fibroblast line **(Fig. 2b-g)**. We confirmed these findings in hESC that also exhibited a similar short G1 and enhanced expression of Cyclin D2 and E (**Supplementary Fig. 2**).

**Figure 2.**
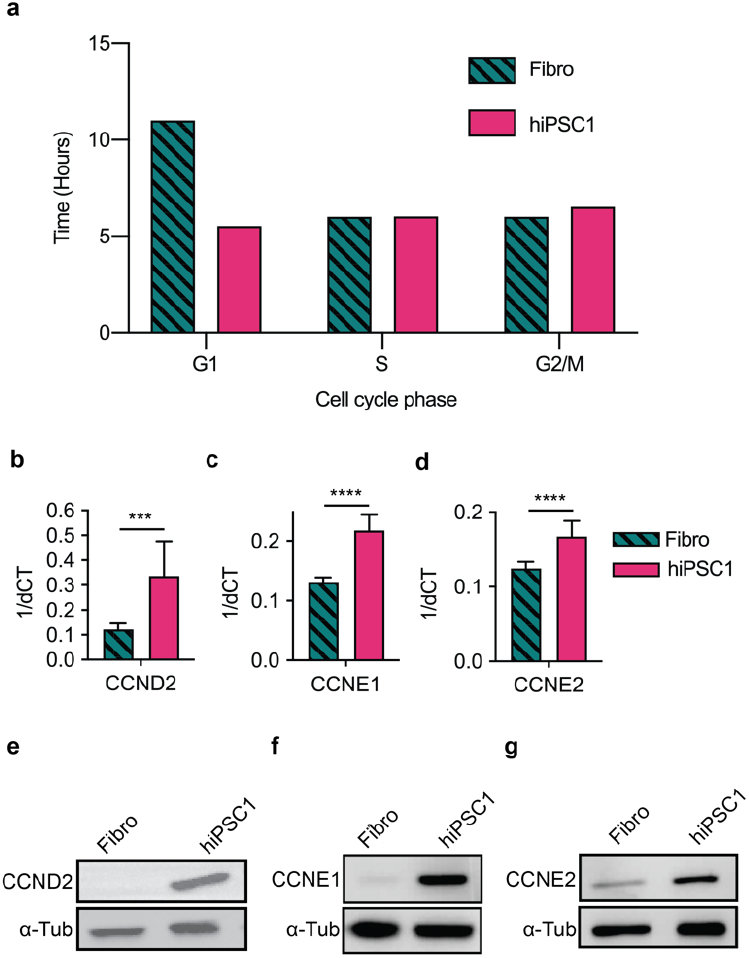
Cell cycle dynamics and cyclin expression in human PSC are candidates for replication stress initiation. a-g, Cell cycle time and cyclin gene expression was compared in isogenic fibroblast (Fibro) and hiPSCI cell lines. **a**, Cell cycle phase time determined from time-lapse microscopy and EdU pulse chase analysis of asynchronous cells. **b-d**, RT-qPCR gene expression data for CCND2 (**b**), CCNE1 (**c**) and CCNE2 (**d**). Data in **b-d**, are mean ± s.d., two-tailed *t*-test, ***P<0.001, ****P<0.0001 *(n* = 3 experiments). t-value; 4.510, 9.546 and 5.619 respectively. **e-g**, Representative western blot of protein expression for CCND2 (**e**), CCNE1 (**f**) and CCNE2 (**g**).

We reasoned that the short G1 may impact on genome damage of PSC, since overexpression of cyclin D2 and E in cancer cells has also been reported to enforce an abbreviation of G1 and consequent replication stress, which can be modulated by exogenous nucleosides(Bester et al., 2011, Takano et al., 2000). When we cultured the PSC in medium containing additional exogenous nucleosides, we observed an increase in DNA fibre lengths and replication fork speed to levels comparable with those observed in the somatic cells **(Fig. 3a-d and Supplementary Fig. 3a-d compared to Fig. 1d-g)**. In addition, we noted fewer CldU-only tracts, indicating fewer forks stalled prior to the addition of the second thymidine analogue, IdU **(Fig. 3e)**. There was also a decrease in replication origin density, with ori-ori distances comparable to those observed in the somatic cells **(Fig. 3f and Supplementary Fig. 3e compared to Fig. 1h**), suggesting that, as a consequence of slower fork speed, the cells were firing from dormant origins in the absence of exogenous nucleosides. Under these conditions we observed a marked decrease in the frequency of DSB in human PSC as indicated by a reduction in the number of cells with more than eight gH2AX foci in S and G2 phase cells upon addition of nucleosides **(Fig. 3g and Supplementary Fig. 3f)** and a decrease in tail moment measured using the neutral comet assay **(Fig. 3h)**. Overall, these results indicate that susceptibility to DNA damage observed in the undifferentiated PSC is a consequence of replication stress that can be alleviated by exogenous nucleosides.

**Figure 3.**
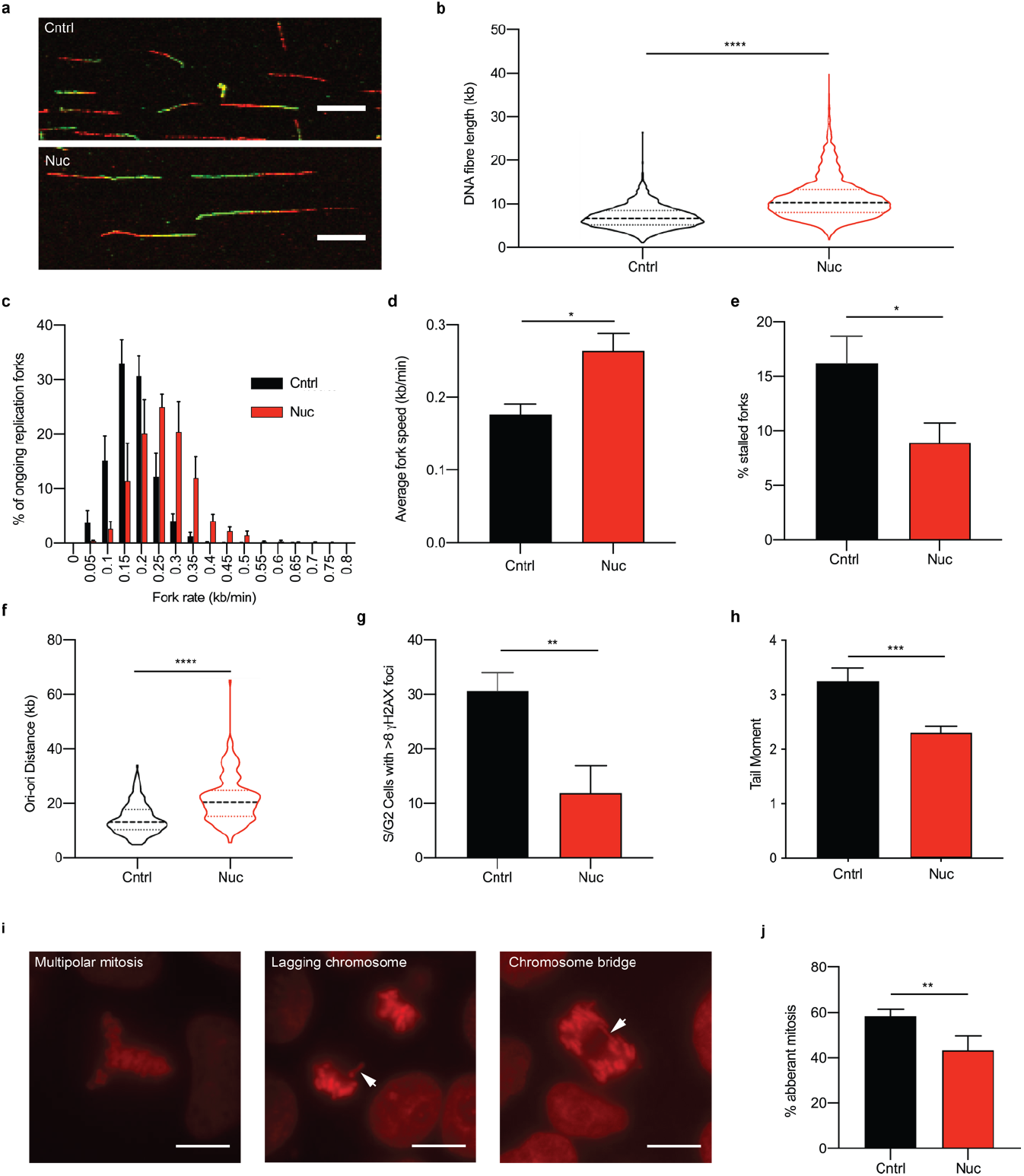
Exogenously supplied nucleosides alleviates replication stress in human PSC. a-e, DNA fibre assays to determine DNA replication dynamics of cells grown in presence of nucleosides (Nuc) and a negative control (Cntrl). **a**, Representative images of DNA fibres in Cntrl (top) and Nuc (bottom) conditions. Scale bar, 10μm. **b** Combined length of CldU and IdU in individual fibres (*n* > 200 forks per cell line per experiment, *n* = 3 experiments). Median distance, 25^th^ and 75^th^ quartiles are presented, two-tailed *t*-test, ***P<0.001. t-value, 40.23. **c**, Distribution of replication fork rates (*n* > 200 forks per condition per experiment, *n* = 3 experiments), data is mean ± s.e.m. **d**, Mean fork rates from (**c**) mean ± s.d., two-tailed *t*-test, *P<0.05 (*n* = 3 experiments). t-value, 3.078. **e**, Frequency of CldU only tracts that denote a stalled replication fork, (*n* > 700 forks per condition per experiment, *n* = 3 experiments) mean ± s.d., two-tailed *t*-test, *P<0.05. t-value, 4.162. **f**, Distribution of adjacent origins distance measurements (Ori-ori). Median distance, 25^th^ and 75^th^ quartiles are presented, two-tailed *t*-test, ****P<0.0001 (*n* > 150 per cell line, *n* = 3 experiments). t-value, 9.815. **g**, Frequency of S/G2 (determined from DNA content) cells with >8 gH2AX foci per cell. Data are mean ± s.d., two-tailed *t*-test, **P<0.01, t-value, 5.333 (*n* > 100 cells per condition per experiment, *n* = 3 experiments). **h**, Average tail moment from neutral comet assay experiments. Median distance, 25^th^ and 75^th^ quartiles are presented, two-tailed *t*-test, ***P<0.001, t-value, 3.322 (*n* ≥ 300 cells per condition per experiment, *n* = 3 experiments). **i, j**, Mitotic errors observed from fluorescently labelled chromatin (histone H2B-RFP). **i**, Representative images of mitotic errors. White arrows point to mitotic error in each case. Scale bar, 10μm **j**, Average frequency of mitotic errors observed. Data are mean ± s.d., unpaired *t*-test, P<0.001, t-value, 4.217 (n = 13-43 mitosis assessed per condition per experiment, *n* = 4 experiments).

A detrimental consequence of replication stress is the presence of under-replicated regions that can persist into mitosis and hinder chromosome separation (Burrell et al., 2013, Bester et al., 2011, Ichijima et al., 2010). This in turn can lead to mitotic aberrations including chromosome bridges, lagging chromosomes and the formation of micronuclei (Janssen et al., 2011, Crasta et al., 2012, Ichijima et al., 2010). Using time-lapse microscopy of hiPSC1 cells stably transfected with H2B-RFP to fluorescently-label chromatin, we tracked the progression of cells through mitosis. Consistent with previous reports (Zhang et al., 2019, Lamm et al., 2016), we observed a high incidence of mitotic errors in human PSC. However, the incidence of these errors was significantly decreased in cells cultured in the presence of nucleosides **(Fig. 3i,j)**, indicating that replication stress in human PSC is a cause of mitotic errors.

To investigate the consequences of these observations for the proliferation of human PSC, we used time-lapse microscopy to track the growth of single cells through successive divisions. When hiPSC1 cells were seeded at low density, 68% of those that attached went on to divide in normal culture medium, whereas 79% entered mitosis in medium supplemented with nucleosides. This is consistent with our previous observation that human PSC activate apoptosis in response to replication stress (Desmarais et al., 2012, Desmarais et al., 2016). Of those that did enter mitosis, 59% went on to form colonies of two or more cells under standard culture conditions, with many cells dying after the first and subsequent divisions (**Fig. 4a and Supplementary Fig. 4a**). By contrast, fewer cells died following division when exogenous nucleosides were added and 91% cells went on to form colonies (**Fig. 4a Supplementary Fig. 4b**) with substantially greater final size (**Fig. 4b**). It is notable that, in the absence of added nucleosides, there was a consistently higher number of abortive cell divisions involving the death of both daughter cells (**Fig. 4c,d**), a result that would be anticipated if the mitotic errors caused by DNA replication stress are catastrophic for both daughter cells.

**Figure 4.**
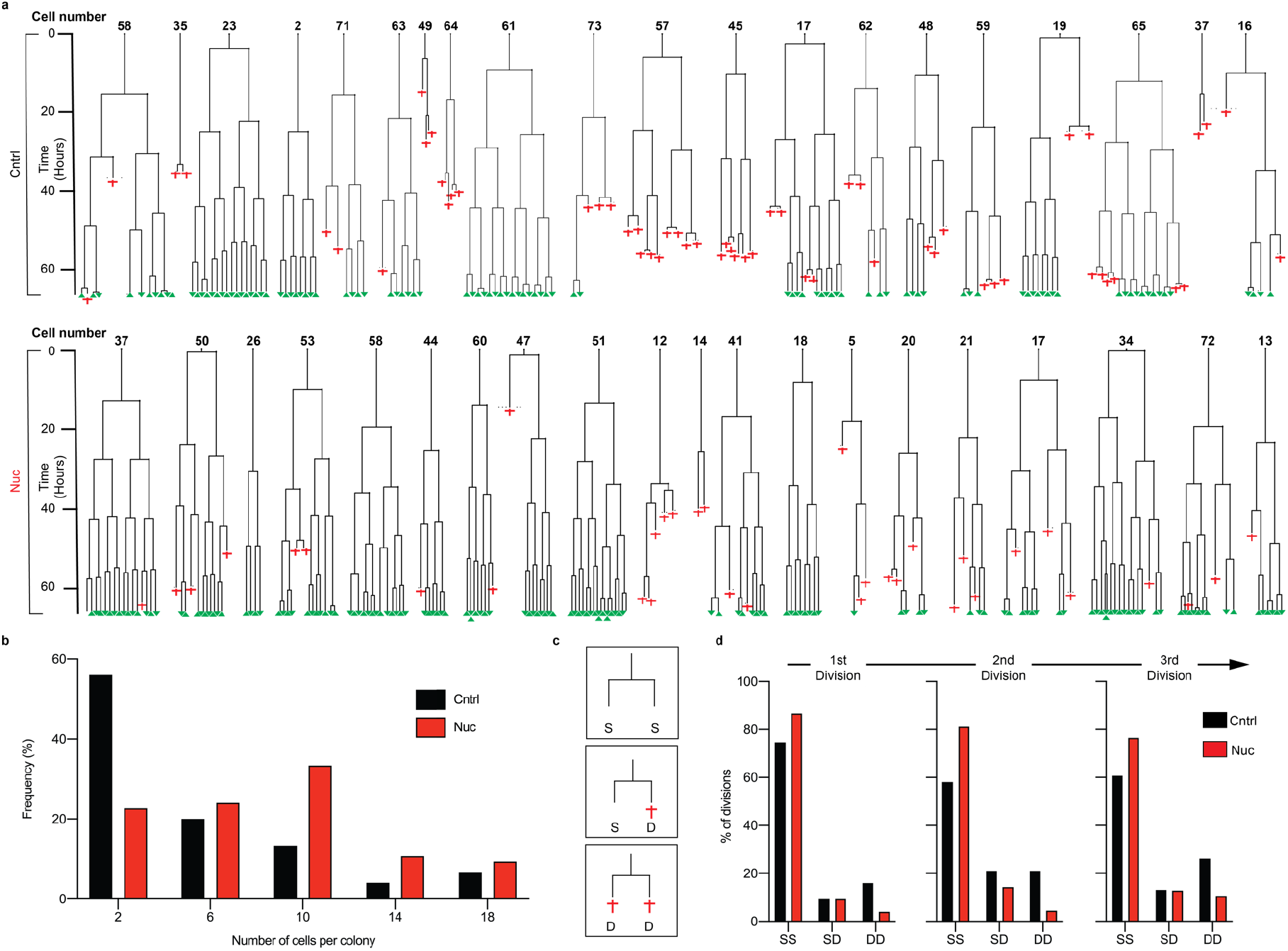
Exogenous nucleosides improve survival and the ability of human PSC to re-enter the cell cycle post plating. a, Time-lapse data tracing the growth of individual hiPSC1 cells that reach the first cell division plotted as lineage trees. A random sample of 20 cells (full set of 75 cells shown in Supplementary Fig. 4) grown in Cntrl (top) and Nuc (bottom) conditions are displayed. Where the trees fork indicates a cell division, red crosses show cell death and surviving cells at the end of the time-lapse are noted with a green triangle. Time is displayed on the y-axis. **b-d**, Summary data of the lineage tree analysis **b**, Histogram of the distribution colony sizes at the end of the time-lapse experiment **c**, Schematic illustrating the scoring method used in **(d).** After each division the fate of the daughter cells was recorded; Both daughter cells surviving (SS), one daughter cell survives and one dies (SD) or both daughter cells die (DD). **d**, Individual bar charts show the frequency of SS, SD and DD daughter cell fates following the 1^st^, 2^nd^ and 3^rd^ divisions (left to right) after plating. Cntrl (black) and Nuc (red) conditions are shown.

Taken together, our results demonstrate that human PSC, compared to somatic cells, are predisposed to high levels of replication stress, manifest by slower rates of DNA synthesis, activation of latent origins of replication and the stalling of replication forks. One consequence is their susceptibility to double stranded DNA breaks that, in turn, may lead to genomic rearrangements during mitosis (Janssen et al., 2011, Ichijima et al., 2010). However, a further feature of human PSC is that, unlike somatic cells, they tend to undergo apoptosis in response to replication stress, so minimising the appearance of mutant cells. This might reflect the demands of cell proliferation in the early embryo in which any genomic damage in even one cell could be catastrophic for the whole embryo (Desmarais et al., 2012, Desmarais et al., 2016). Indeed, in a separate study we have found that the overall mutation rate in human PSC is rather low (Thompson et al., 2019), despite their propensity to DNA damage. That human PSC, nevertheless, tend to accumulate particular recurrent mutations and genomic rearrangements most likely reflects selection for growth advantages among those few variants that escape apoptosis. It is notable that resistance to apoptosis is a common feature of the recurrent variants that do arise in human PSC (Merkle et al., 2017, Avery et al., 2013). Our observation that exogenous nucleosides substantially reduced DNA replication stress in human PSC, perhaps compensating for metabolic changes that stem from their shortened G1 and relaxed G1/S transition, without disrupting pluripotency (**Supplementary Fig. 5–6**), provides a means to reduce the incidence of recurrent genetic changes that may compromise the use of human PSC for disease modelling and regenerative medicine.

**Supplementary Figure 1.**
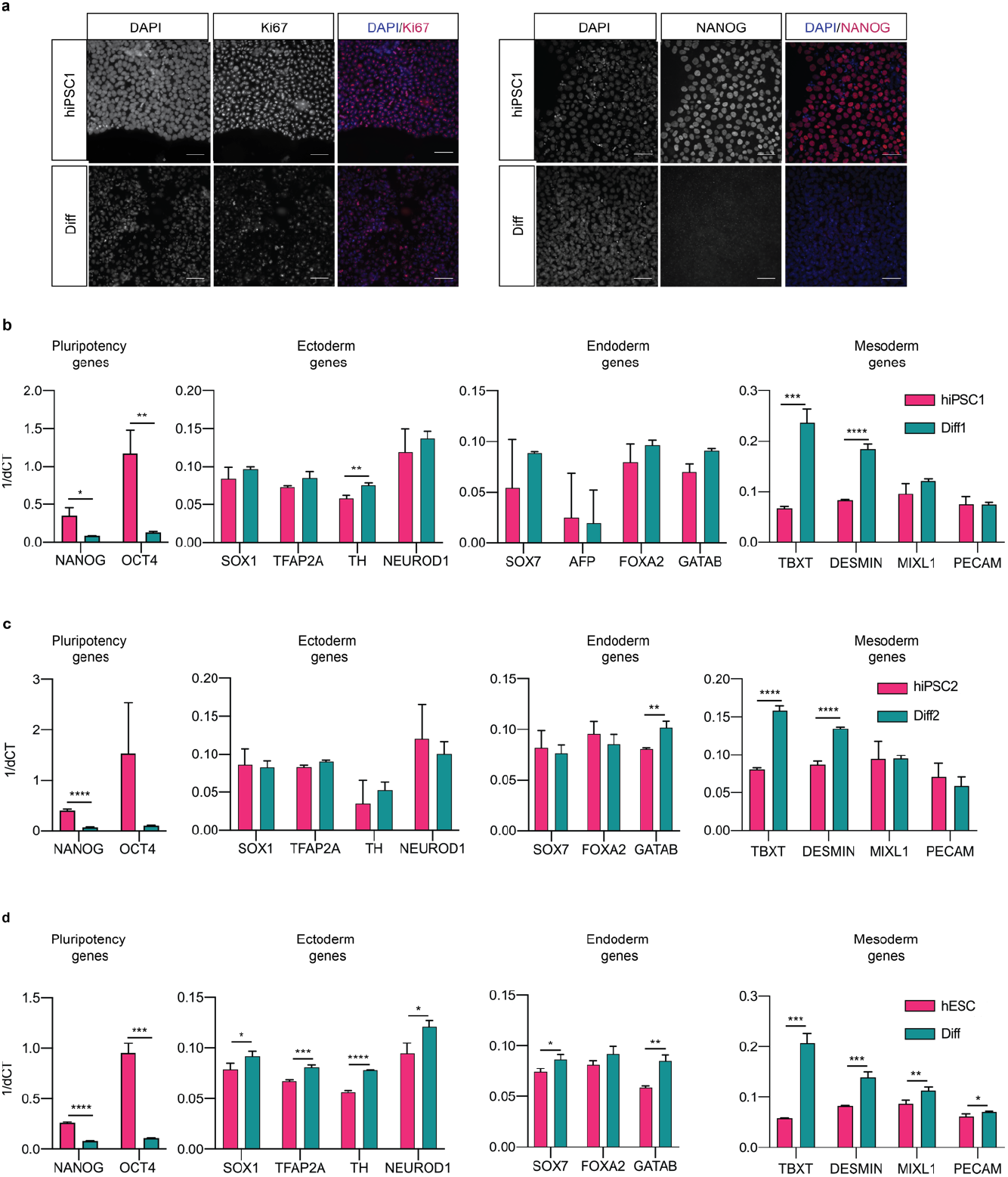
Differentiation of human PSC lines to obtain isogenic differentiated derivatives. a, Representative immunofluorescence images of hiPSC1 and hiPSC Differentiated cells stained for Ki67 and NANOG, with nuclei counterstained with Hoechst 33342. Scale bar, 50μm. **b-d**, RT-qPCR gene expression data of hiPSC1 **(a)**, hiPSC2 **(b)** and hESC **(c)** compared to their differentiated derivatives. Genes associated with pluripotency, ectoderm, endoderm and mesoderm are displayed (left to right). Data in **b**-**d**, are mean ± s.d., two-tailed *t*-test, *P<0.05, **P<0.01, ***P<0.001, ****P<0.0001, (*n* = 3 experiments).

**Supplementary Figure 2.**
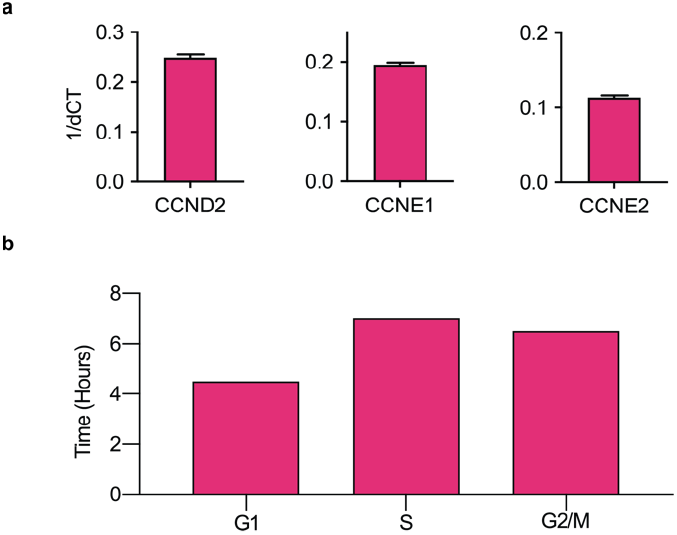
Shortened G1 phase and high G1 cyclin expression in multiple PSC lines. a, RT-qPCR gene expression data for *CCND2*, *CCNE1* and *CCNE2* in hESC line **b**, The cell cycle phase time, determined from time-lapse microscopy and EdU pulse chase analysis in hESC line.

**Supplementary Figure 3.**
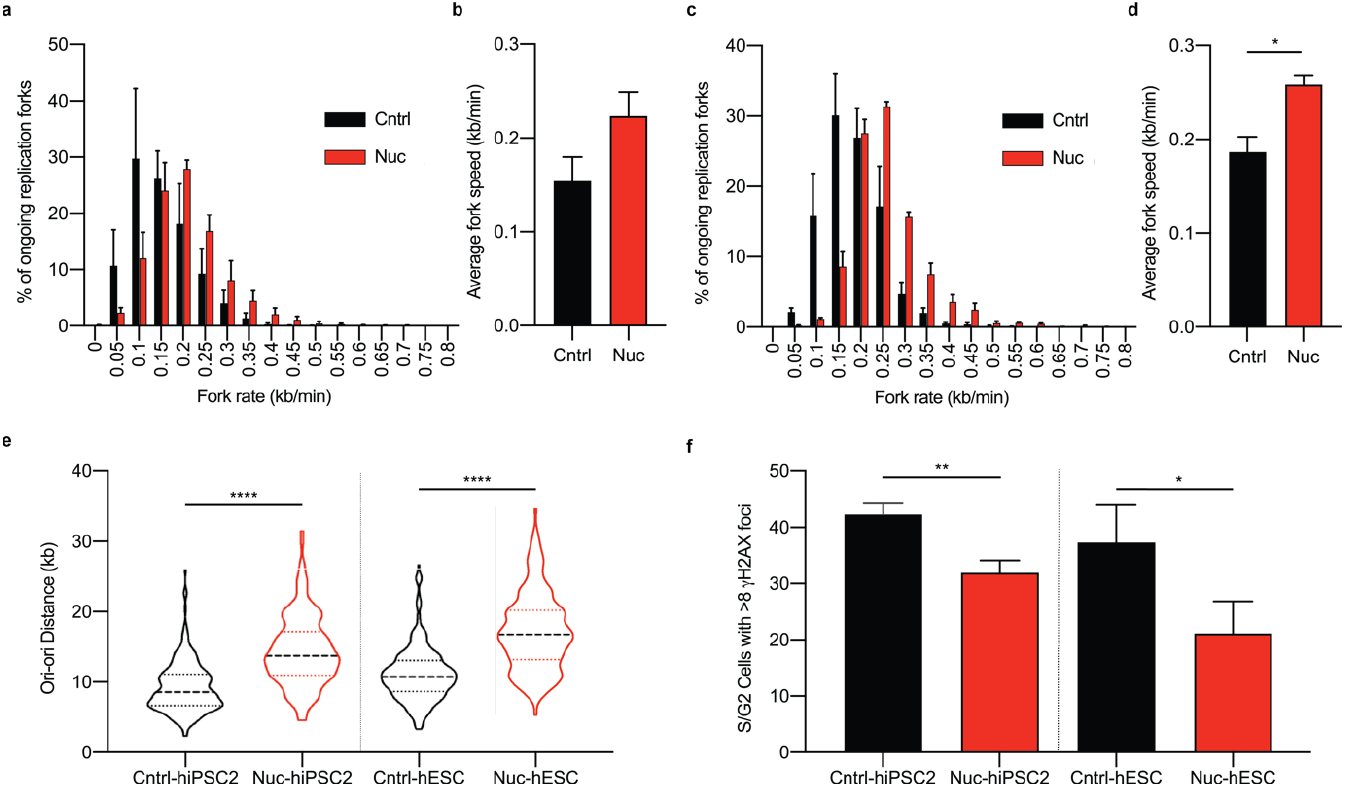
Replication stress and a susceptibility to DNA damage is a characteristic of undifferentiated human PSC, additional cell lines. a-e, Data presented from DNA fibre assays; replication rates and replication origin distances from individually assessed replication forks. Comparison of replication fork rates made between control (Cntrl) and Nucleoside (Nuc) conditions for hiPSC2 **(a,b)** and hESC **(c,d). a,c** Distribution of replication fork rates (*n* > 200 forks per cell line per experiment, *n* = 3 experiments), data presented is the mean value from each experiment ± s.e.m. **b,d** Mean fork rates. Data is mean ± s.d., two-tailed *t*-test, *P<0.05. (*n* > 200 forks per cell line per experiment, *n* = 3 experiments) **e**, Distribution of adjacent origins distance measurements (Ori-ori). Median distance, 25^th^ and 75^th^ quartiles are presented, two-tailed *t*-test, ****P<0.0001 (*n* > 160 per cell line, *n* = 3 experiments). **f**, Double strand breaks assessed by counting the number of cells with ≥8 gH2AX foci in S/G2 phase cells determined from DNA content. Comparison made between hiPSC2 and hESC cells grown in control (Cntrl) and nucleoside (Nuc) conditions. Data in **f** are mean values of 3 independent experiments ± s.d., two-tailed *t*-test, *P<0.05, **P<0.01 (*n* > 100 cells per cell line per experiment).

**Supplementary Figure 4.**
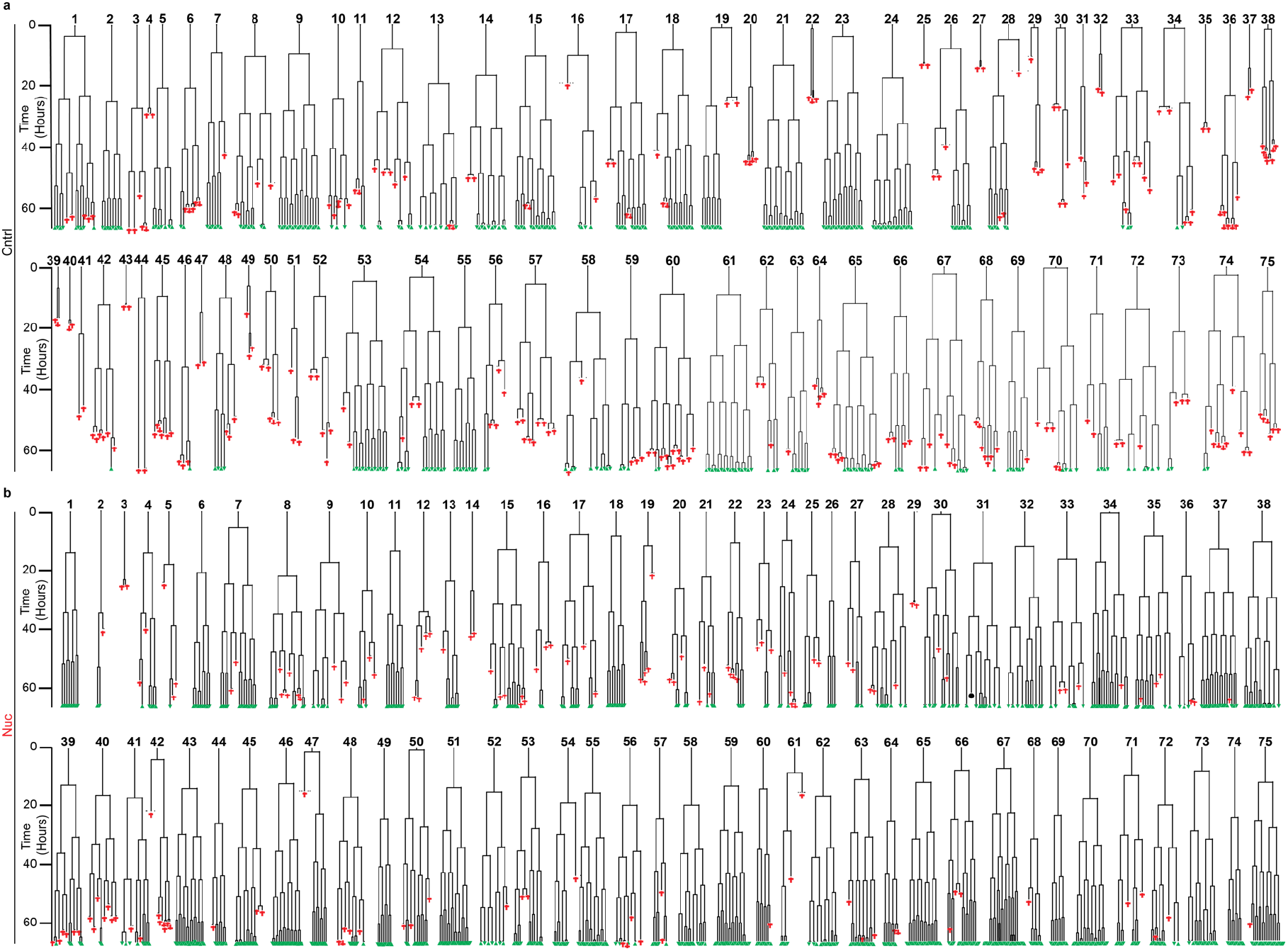
Exogenous nucleosides improve survival and plating efficiency of human PSC. a,b, Lineage trees of the 75 cells, that reached the first cell division, sampled from Cntrl (**a**) and Nuc (**b**) conditions. 20 randomly selected starting cells and the resulting lineage trees are displayed in Figure 4.

**Supplementary Figure 5.**
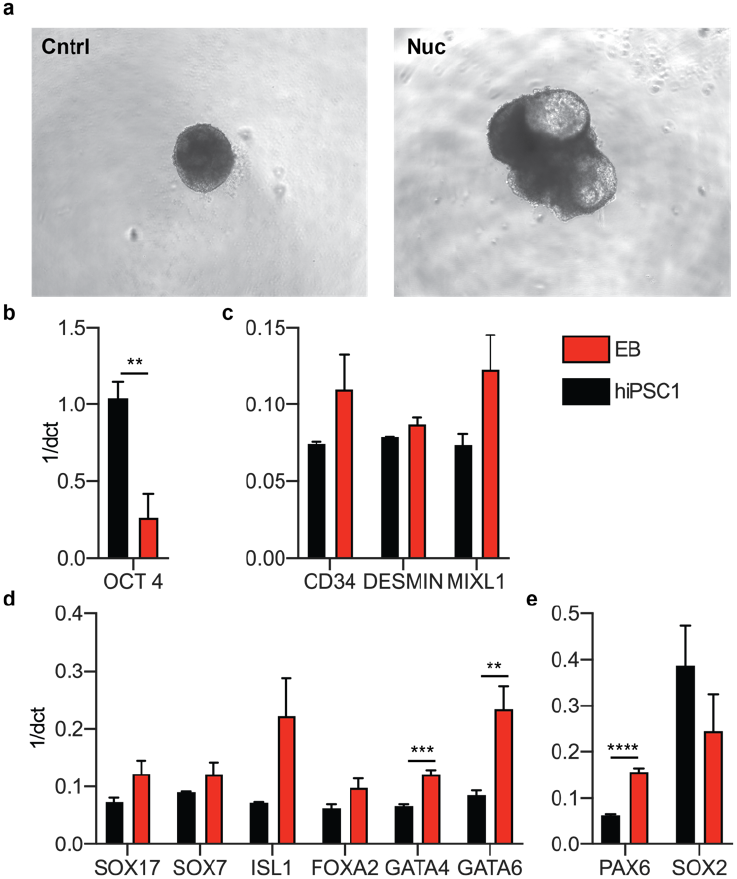
Human PSC retain the ability to differentiate into the three germ layers when cultured in exogenous nucleosides a, Representative images of 10-day embryoid bodies. Prior to differentiation the hiPSC1 cells were expanded in control (Cntrl) or with exogenous nucleosides (Nuc) for ten passages. **b-d**, RT-qPCR gene expression data of hiPSC1 compared to 10-day embryoid bodies. Genes associated with pluripotency **(b)**, mesoderm **(c)**, endoderm **(d)** and ectoderm **(e)** are displayed. Data in **b**-**d**, are mean ± s.d., two-tailed *t*-test, **P<0.01, ***P<0.001, ****P<0.0001 (*n* = 3 experiments).

**Supplementary Figure 6.**
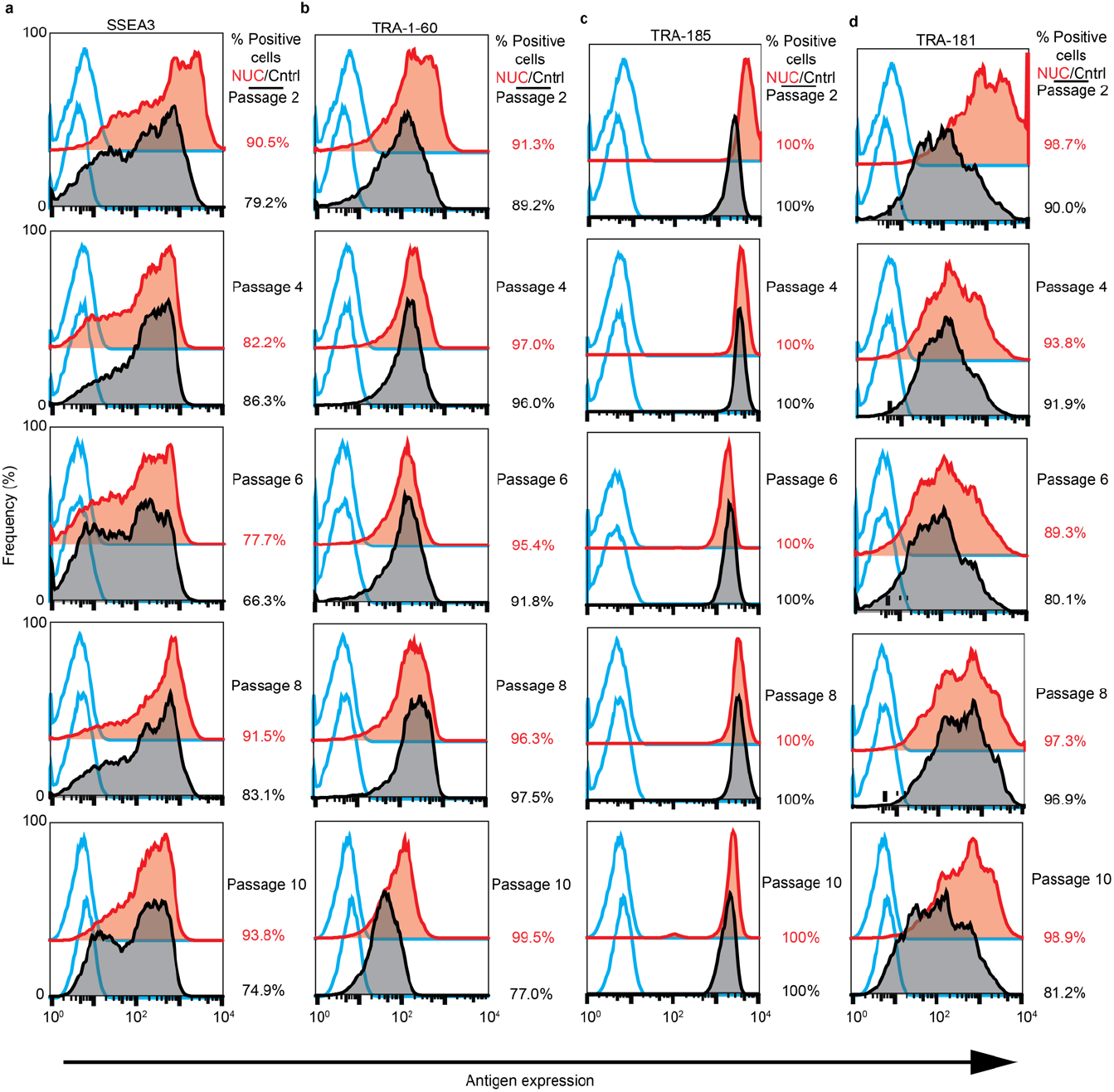
Exogenous nucleosides do not alter the pluripotency associated antigen expression. a-d, Pluripotency associated antigen expression, human PSC are grown with exogenous nucleosides (NUC) (Red) versus a control (Cntrl) (Black) over 10 passages. Baseline fluorescence was set using the primary antibody control, P3X (Blue). Percentage positive population is displayed to the right of each histogram. **a**, SSEA3 antigen expression from passage 2 to passage 10 (top to bottom). **b**, TRA-1-60 antigen expression passage 2 to 10. **c**, Pluripotency associated antigen expression of TRA-185 cultured for 10 passages. **d**, Tra-181 antigen expression from passage 2 to passage 10. **a-d**, Modal scaled channels, histograms are displayed as a percentage of the maximum count.

## Methods

### Human pluripotent stem cell culture

Human embryonic stem cells (hESCs) and human induced pluripotent stem cells (hiPSCs) were cultured on Vitronectin (VTN-N) recombinant human protein (ThermoFisher Scientific, A14700). Culture vessels were coated with 200μL/cm^2^ of Vitronectin that had been diluted to 6μg/ml with PBS and incubated at 37°C for at least 1 hour. The cells were maintained in feeder free conditions, batch fed daily with Essential 8(Chen et al., 2011) or mTeSR^⅄^1 (STEMCELL Technologies, 85850) cell culture media, and incubated in a 37°C, 5% CO_2_ humidified incubator. Passaging of cells was performed by clump dissociation using ReLeSR (STEMCELL technologies, 05873) following manufacturer guidelines.

### Differentiation of human PSC

Human PSC were grown for 5 days in E8 media(Chen et al., 2011) without FGF-2 and TGF-b but supplemented with 10μM CHIR99021 (Tocris, 4423). Loss of pluripotency was confirmed by RT-qPCR panel of self-renewal, mesoderm, endoderm and ectoderm genes (Table 1) and by immunofluorescence staining and imaging of NANOG.

**Table 1.**
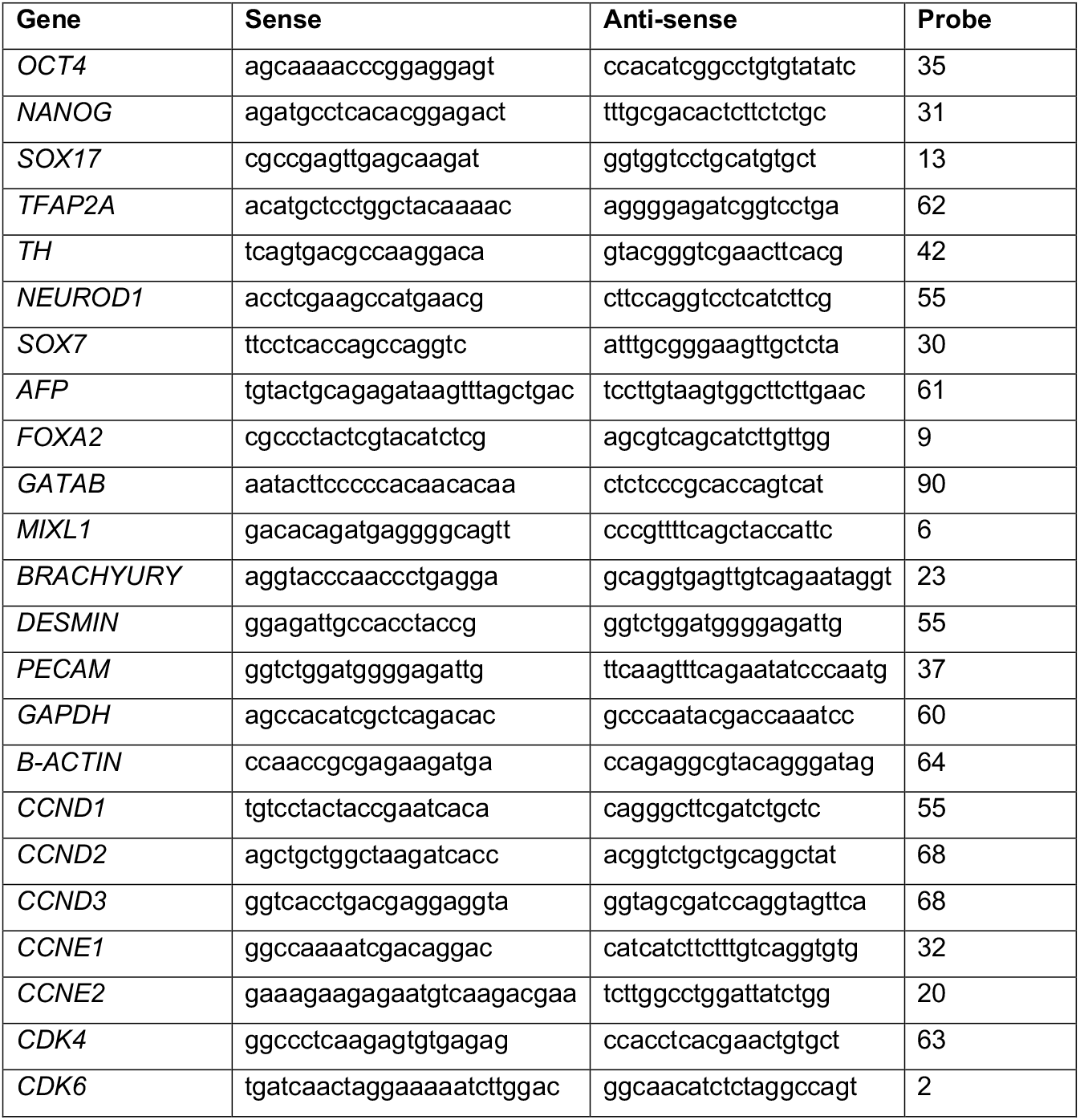
Primer sequences and Universal probe library probes.

### Fibroblast cell culture

Fibroblasts (ATCC, CRL2429) were grown in Iscove’s Modified Dulbecco’s Medium (Thermo Fisher Scientific, 12440053) with 20% FBS (HyCLone, SV30160.03). Cells were passaged using TrypLE cell dissociation enzyme (Thermo Fisher Scientific, 12504013) following manufacturer’s guidelines. Cells were maintained at 37°C and 5% CO_2_ in a humidified incubator.

### Nucleoside supplementation

Embryomax Nucleosides 100X (Merck, ES-008-D) were added to mTeSR cell culture media at a final concentration of 0.5X. All experiments were performed after 72 hours in culture with the supplementation of nucleosides.

### Immunocytochemistry

Cells were fixed with 4% paraformaldehyde (Sigma, 158127) for 10 minutes. Cells were permeabilised and blocked with 0.3% Triton-X (Sigma, T8787), 10% goat serum (Thermo Fisher Scientific, 16210072) and 3% BSA (Sigma) in PBS for 1 hour. Primary antibody incubation was performed overnight at 4°C: Anti-Phospho-Histone H2A.X (Ser139) (Cell Signalling Technologies, 9718; 1:400), Anti-gamma H2A.X (Phospho S139) (abcam, ab26350; 1:500), Anti-Nanog (Cell Signalling, 4903; diluted 1:500), Anti-Nanog (Cell Signalling, 4893; diluted 1:500) and Anti-Ki67 (abcam, ab238020; diluted 1:100). Secondary antibody incubation was performed for 1 hour: Alexa Fluor 488-conjugated anti-rabbit IgG(Life Technologies, A11034; diluted 1:400), Alexa Fluor 647 AffiniPure Goat anti-Rabbit IgG (H+L) (Jackson Immuno Research, 111-605-003; 1:1000) and Alexa Fluor 647 AffiniPure Goat anti-Mouse IgG (H+L) (Jackson Immuno Research, 115-605-003; 1:400). All antibodies were diluted in 1% BSA (Sigma) and 0.3% Triton-X (Sigma, T8787) in PBS. Nuclei were counterstained with Hoechst 33342 (Thermo Fisher Scientific, H3570; diluted 1:1000) and images were acquired using an INCell Analyzer 2200 (GE Healthcare) high content microscope taking 25 or 30 randomized images per well.

### Immunofluorescence data analysis

Where possible, CellProfiler(Carpenter et al., 2006) cell image analysis software was used to analyse the high content images.

Immunofluorescence was quantified above a threshold set by a secondary only control. Hoechst 33342 was used to identify the cells nuclei and mask the nuclear area over the immunofluorescence staining. Where it was necessary to separate cells by cell cycle stage the integrated intensity of the stained nuclei was calculated using CellProfiler Analyst(Jones et al., 2008).

### Neutral comet assay

150μL of 0.6% agarose (Sigma, A9539) was set on a fully frosted glass slide, sandwiched beneath a coverslip. Once dried, 12,000 cells per conditions was resuspended in 75μL ice cold PBS and mixed with 75μL of 1.2% low melting agarose (Sigma, A4018). The cell and agarose suspension was mounted on top of the original agarose layer beneath a coverslip and set at 4°C. The slides were immersed in pre-chilled lysis buffer (2.5M NaCl, 10mM Tris-HCL, 100mM EDTA pH8.0, 0.5% Triton-X, 3% DMSO) for 1.5 hours at 4°C, washed in H_2_O and equilibrized in electrophoresis buffer (300mM sodium acetate, 100mM Tris-EDTA and 1% DMSO) for 1 hour. Electrophoresis is performed at 25V for 1 hour. Slides were stained with SYBR green (Sigma, S9430), imaged and quantified using Comet Assay IV (Instem) live video measurement system.

### DNA fibre assay

DNA fibre assay was performed as previously described(Groth et al., 2010). Briefly, cells were plated and grown for a minimum of 72 hours before sequential pulse labelling with 2.5mM CldU (Sigma, C6891; 1:100) and then 2.5mM IdU (Sigma, I7125; 1:10) for 20 minutes each. Cells were washed with ice cold PBS, dissociated using TrypLE cell dissociation enzyme (Thermo Fisher Scientific, 12504013) and diluted to 3.5×10^5^ cells/ml in cold PBS. To spread the labelled fibers, 2μL of cell suspension was dropped onto a glass slide and allowed to dry for 5-7 minutes before adding 7μL of spreading buffer (200mM Tris-HCL PH7.4, 50mM EDTA, 0.5% SDS). The solutions were mixed with a pipette tip and incubated for 2 minutes. Slides were tilted at an angle of 10° and the droplet timed to ensure consistent spreading was achieved. Slides were air dried and fixed with 3:1 methanol/acetic acid. For the immunostaining, the glass slides were first washed twice with H2O for 5 minutes each, denatured with 2.5M HCL for 1 hour and then blocked in 1% BSA (Sigma) and 0.1% Tween20. Primary antibodies were incubated for 1 hour: Rat anti-BrdU, clone BU1/75 (Novus Biologicals NB500-169) (AbD Serotec; diluted 1:400) or Anti-BrdU clone BU1/75 (ICR1) (abcam, ab6326; diluted 1:400) and Mouse anti-BrDU (Clone B44) (Becton Dickinson, 347580; diluted 1:250). The secondary antibodies used were Alexa Fluor 555 goat anti-rat IgG (Thermo Fisher Scientific, A21434; diluted 1:500) and Alexa Fluor 488 F (ab’)2-Goat anti-Mouse IgG (Thermo Fisher Scientific, A-11017; diluted 1:500). Slides were mounted with Fluoroshield (Sigma, F6182), and images were acquired using Olympus FV1000 confocal microscope.

### Cell cycle time analysis

Total cell cycle time was measured from the time elapsed between cells first and second divisions monitored by time-lapse microscopy. Cells were seeded at 500 cells/cm^2^ and images were acquired every 10 minutes for 48-72 hours on a Nikon Biostation CT with a 20X objective lens. Images were compiled in CL Quant (NIKON) and analysed using FIJI (ImageJ).

Individual phase times were measured using previously reported pulse chase analysis(Begg et al., 1985). Briefly, cells were seeded into multi-well plates and pulse labelled with 10μM EdU for 45 minutes (Click-iT EdU Alexa Fluor 647 Flow Cytometry Assay Kit, Thermo Fisher Scientific, C10424). At hourly intervals for 24-32 hours a single well was harvested using TrypLE cell dissociation enzyme (Thermo Fisher Scientific, 12504013), pelleted and fixed using Click-iT fixative. After washing with 10% FCS and 1% BSA the cells were permeabilised with the Click-iT saponin based wash buffer before staining with the Click-iT reaction cocktail and counterstaining with Hoechst 33342 (Thermo Fisher Scientific, H3570; diluted 1:1000). A minimum of 10,000 dual labelled cells were recorded by FACs cytometry. Relative movement and mid-S phase movement was calculated using FLOWJO single-cell flow cytometry analysis software (Becton Dickinson).

### RNA Extraction and reverse transcriptase qPCR

RNA was extracted using Qiagen RNeasy kit. cDNA synthesis was performed using high capacity reverse transcription kit (Thermo Fisher Scientific, 4368814). qPCR was performed in 384 well plates with 10μL reactions consisting of 1X TaqMan Fast Universal Master Mix (ThermoFisher, 4352042), 100nM of forward and reverse primers (Table 1), 100nM of probe from the Universal Probe Library (Roche) and 2μL of 5ng/μL cDNA. PCR reactions were run on the QuantStudio 12K Flex Thermocycler (Life Technologies 4471087). All reactions were performed in triplicate with comparative Ct normalized to GAPDH or B-ACTIN expression.

## Western blotting

Laemili buffer (4% SDS, 20% Glycerol, 0.125M Tris HCl, 0.004% bromphenol blue) was added to cell pellets and sonicated for 10 seconds. Protein lysate was incubated for 10 minutes at 95°C and concentration was determined by NanoDrop spectrophotometer (Thermo Fisher Scientific).

Protein was separated on 10% ProtoGel (National Diagnostics) run at 120V for 1.5 hours and transferred onto PVDF membrane (Millipore, #IPVH00010). Primary antibodies were incubated over night at 4°C: α-Tubulin (Cell Signalling Technology, 2144; diluted 1:1000),

Cyclin E1 (D7T3U) (Cell Signalling Technology, 20808; diluted 1:500), Cyclin E2 (Cell Signalling Technology, 4132; diluted 1:500), Cyclin D2 (D52F9) (Cell Signalling Technology, 3741; diluted 1:500). The blot was washed and incubated with Anti-rabbit IgG or anti-mouse IgG secondary antibody for 1 hour (Promega, W401 & W402). Immunoreactivity was visualised with ECL prime (GE Healthcare, RPN2232) on a CCD-based camera.

## The generation of hPSC stably expressing H2B-RFP

Transfection with pCAG-H2B-RFP-IRES-PURO vector(Liew et al., 2007) was performed using electroporation. Cells were electroporated with a single 1600V pulse for 20msec using the Neon transfection system according to manufacturer’s instructions (Thermo Fisher Scientific, MPK10025). Stable clones were obtained by puromycin selection. Puromycin concentration was increased gradually over 5 days to a final concentration of 0.375μg/mL before flow sorting for the brightest population of RFP.

The data sets used to generate the figures in this article are available upon reasonable request from the corresponding author.

## Author Contributions

Peter Andrews and Ivana Barbaric oversaw the project. Jason Halliwell, Ivana Barbaric and Peter Andrews devised the experiments. Jason Halliwell performed most cell biology experiments with help from Thomas Frith, Owen Laing, Christopher Price, Dylan Stavish and Oliver Bower. Jason Halliwell and Thomas Frith performed the differentiations of the human PSC. Jason Halliwell and Owen Laing performed embryoid body formation and pluripotency associated antigen expression analysis. Jason Halliwell, Christopher Price, Dylan Stavish and Oliver Bower performed time-lapse lineage tree experimentation and analysis. Zoe Hewitt oversaw the derivation of the hiPSC2 cell line. Peter Andrews, Ivana Barbaric, Paul Gokhale and Sherif El-Khamisy provided experimental advice.

The manuscript was drafted by Jason Halliwell, Peter Andrews and Ivana barbaric.

## Acknowledgements

This project has received funding from the European Union’s Horizon 2020 research and innovation programme under grant agreement No. 668724.

This work was partly funded by the European Union’s Horizon 2020 research and innovation program under grant agreement No. 668724 and partly by the UK Regenerative Medicine Platform, MRC reference MR/R015724/1.

## References

Ahuja, A. K., Jodkowska, K., Teloni, F., Bizard, A. H., Zellweger, R., Herrador, R., Ortega, S., Hickson, I. D., Altmeyer, M., Mendez, J. & Lopes, M. 2016. A short G1 phase imposes constitutive replication stress and fork remodelling in mouse embryonic stem cells. Nat Commun, 7, 10660.

Amps, K., Andrews, P. W., Anyfantis, G., Armstrong, L., Avery, S., Baharvand, H., Baker, J., Baker, D., Munoz, M. B., Beil, S., Benvenisty, N., Ben-Yosef, D., Biancotti, J. C., Bosman, A., Brena, R. M., Brison, D., Caisander, G., Camarasa, M. V., Chen, J., Chiao, E., Choi, Y. M., Choo, A. B., Collins, D., Colman, A., Crook, J. M., Daley, G. Q., Dalton, A., De Sousa, P. A., Denning, C., Downie, J., Dvorak, P., Montgomery, K. D., Feki, A., Ford, A., Fox, V., Fraga, A. M., Frumkin, T., Ge, L., Gokhale, P. J., Golan-Lev, T., Gourabi, H., Gropp, M., Lu, G., Hampl, A., Harron, K., Healy, L., Herath, W., Holm, F., Hovatta, O., Hyllner, J., Inamdar, M. S., Irwanto, A. K., Ishii, T., Jaconi, M., Jin, Y., Kimber, S., Kiselev, S., Knowles, B. B., Kopper, O., Kukharenko, V., Kuliev, A., Lagarkova, M. A., Laird, P. W., Lako, M., Laslett, A. L., Lavon, N., Lee, D. R., Lee, J. E., Li, C., Lim, L. S., Ludwig, T. E., Ma, Y., Maltby, E., Mateizel, I., Mayshar, Y., Mileikovsky, M., Minger, S. L., Miyazaki, T., Moon, S. Y., Moore, H., Mummery, C., Nagy, A., Nakatsuji, N., Narwani, K., Oh, S. K., Olson, C., Otonkoski, T., Pan, F., Park, I. H., Pells, S., Pera, M. F., Pereira, L. V., Qi, O., Raj, G. S., Reubinoff, B., Robins, A., Robson, P., Rossant, J., Salekdeh, G. H., Schulz, T. C., et al. 2011. Screening ethnically diverse human embryonic stem cells identifies a chromosome 20 minimal amplicon conferring growth advantage. Nat Biotechnol, 29, 1132–44.

Avery, S., Hirst, A. J., Baker, D., Lim, C. Y., Alagaratnam, S., Skotheim, R. I., Lothe, R. A., Pera, M. F., Colman, A., Robson, P., Andrews, P. W. & Knowles, B. B. 2013. BCL-XL mediates the strong selective advantage of a 20q11.21 amplification commonly found in human embryonic stem cell cultures. Stem Cell Reports, 1, 379–86.

Becker, K. A., Ghule, P. N., Lian, J. B., Stein, J. L., Van Wijnen, A. J. & Stein, G. S. 2010. Cyclin D2 and the CDK substrate p220(NPAT) are required for self-renewal of human embryonic stem cells. J Cell Physiol, 222, 456–64.

Becker, K. A., Ghule, P. N., Therrien, J. A., Lian, J. B., Stein, J. L., Van Wijnen, A. J. & Stein, G. S. 2006. Self-renewal of human embryonic stem cells is supported by a shortened G1 cell cycle phase. J Cell Physiol, 209, 883–93.

Begg, A. C., Mcnally, N. J., Shrieve, D. C. & KÄRCHER, H. 1985. A method to measure the duration of DNA synthesis and the potential doubling time from a single sample. Cytometry, 6, 620–6.

Bester, A. C., Roniger, M., Oren, Y. S., Im, M. M., Sarni, D., Chaoat, M., Bensimon, A., Zamir, G., Shewach, D. S. & Kerem, B. 2011. Nucleotide deficiency promotes genomic instability in early stages of cancer development. Cell, 145, 435–46.

Burrell, R. A., Mcclelland, S. E., Endesfelder, D., Groth, P., Weller, M. C., Shaikh, N., Domingo, E., Kanu, N., Dewhurst, S. M., Gronroos, E., Chew, S. K., Rowan, A. J., Schenk, A., Sheffer, M., Howell, M., Kschischo, M., Behrens, A., Helleday, T., Bartek, J., Tomlinson, I. P. & Swanton, C. 2013. Replication stress links structural and numerical cancer chromosomal instability. Nature, 494, 492–496.

Carpenter, A. E., Jones, T. R., Lamprecht, M. R., Clarke, C., Kang, I. H., Friman, O., Guertin, D. A., Chang, J. H., Lindquist, R. A., Moffat, J., Golland, P. & Sabatini, D. M. 2006. CellProfiler: image analysis software for identifying and quantifying cell phenotypes. Genome Biol, 7, R100.

Chen, G., Gulbranson, D. R., Hou, Z., Bolin, J. M., Ruotti, V., Probasco, M. D., Smuga-Otto, K., Howden, S. E., Diol, N. R., Propson, N. E., Wagner, R., Lee, G. O., Antosiewicz-Bourget, J., Teng, J. M. & Thomson, J. A. 2011. Chemically defined conditions for human iPSC derivation and culture. Nat Methods, 8, 424–9.

Crasta, K., Ganem, N. J., Dagher, R., Lantermann, A. B., Ivanova, E. V., Pan, Y., Nezi, L., Protopopov, A., Chowdhury, D. & Pellman, D. 2012. DNA breaks and chromosome pulverization from errors in mitosis. Nature, 482, 53–8.

Desmarais, J. A., Hoffmann, M. J., Bingham, G., Gagou, M. E., Meuth, M. & Andrews, P. W. 2012. Human embryonic stem cells fail to activate CHK1 and commit to apoptosis in response to DNA replication stress. Stem Cells, 30, 1385–93.

Desmarais, J. A., Unger, C., Damjanov, I., Meuth, M. & Andrews, P. 2016. Apoptosis and failure of checkpoint kinase 1 activation in human induced pluripotent stem cells under replication stress. Stem Cell Res Ther, 7, 17.

Draper, J. S., Smith, K., Gokhale, P., Moore, H. D., Maltby, E., Johnson, J., Meisner, L., Zwaka, T. P., Thomson, J. A. & Andrews, P. W. 2004. Recurrent gain of chromosomes 17q and 12 in cultured human embryonic stem cells. Nat Biotechnol, 22, 53–4.

Groth, P., AuslÄNDER, S., Majumder, M. M., Schultz, N., Johansson, F., Petermann, E. & Helleday, T. 2010. Methylated DNA causes a physical block to replication forks independently of damage signalling, O(6)-methylguanine or DNA single-strand breaks and results in DNA damage. J Mol Biol, 402, 70–82.

Hinds, P. W., Mittnacht, S., Dulic, V., Arnold, A., Reed, S. I. & Weinberg, R. A. 1992. Regulation of retinoblastoma protein functions by ectopic expression of human cyclins. Cell, 70, 993–1006.

Ichijima, Y., Yoshioka, K., Yoshioka, Y., Shinohe, K., Fujimori, H., Unno, J., Takagi, M., Goto, H., Inagaki, M., Mizutani, S. & Teraoka, H. 2010. DNA lesions induced by replication stress trigger mitotic aberration and tetraploidy development. PLoS One, 5, e8821.

Janssen, A., Van Der Burg, M., Szuhai, K., Kops, G. J. & Medema, R. H. 2011. Chromosome segregation errors as a cause of DNA damage and structural chromosome aberrations. Science, 333, 1895–8.

Jones, T. R., Kang, I. H., Wheeler, D. B., Lindquist, R. A., Papallo, A., Sabatini, D. M., Golland, P. & Carpenter, A. E. 2008. CellProfiler Analyst: data exploration and analysis software for complex image-based screens. BMC Bioinformatics, 9, 482.

Lamm, N., Ben-David, U., Golan-Lev, T., StorchovÁ, Z., Benvenisty, N. & Kerem, B. 2016. Genomic Instability in Human Pluripotent Stem Cells Arises from Replicative Stress and Chromosome Condensation Defects. Cell Stem Cell, 18, 253–61.

Liew, C. G., Draper, J. S., Walsh, J., Moore, H. & Andrews, P. W. 2007. Transient and stable transgene expression in human embryonic stem cells. Stem Cells, 25, 1521–8.

Lundberg, A. S. & Weinberg, R. A. 1998. Functional inactivation of the retinoblastoma protein requires sequential modification by at least two distinct cyclin-cdk complexes. Mol Cell Biol, 18, 753–61.

Merkle, F. T., Ghosh, S., Kamitaki, N., Mitchell, J., Avior, Y., Mello, C., Kashin, S., Mekhoubad, S., Ilic, D., Charlton, M., Saphier, G., Handsaker, R. E., Genovese, G., Bar, S., Benvenisty, N., Mccarroll, S. A. & Eggan, K. 2017. Human pluripotent stem cells recurrently acquire and expand dominant negative P53 mutations. Nature, 545, 229–233.

Olariu, V., Harrison, N. J., Coca, D., Gokhale, P. J., Baker, D., Billings, S., Kadirkamanathan, V. & Andrews, P. W. 2010. Modeling the evolution of culture-adapted human embryonic stem cells. Stem Cell Res, 4, 50–6.

Seth, T., Benjamin, N., Michael, N., Tori, S.-B., Kimberley, L., Erik, M. & Karen, M. 2011. Karyotypic abnormalities in human induced pluripotent stem cells and embryonic stem cells. Nature Biotechnology, 29, 313–314.

Takano, Y., Kato, Y., Van Diest, P. J., Masuda, M., Mitomi, H. & Okayasu, I. 2000. Cyclin D2 overexpression and lack of p27 correlate positively and cyclin E inversely with a poor prognosis in gastric cancer cases. Am J Pathol, 156, 585–94.

Thompson, O., Von Meyenn, F., Hewitt, Z., Alexander, J., Wood, A., Weightman, R., Gregory, S., Krueger, F., Andrews, S., Barbaric, I., Gokhale, P. J., Moore, H. D., Reik, W., Milo, M., Nik-Zainal, S., Yusa, K. & Andrews, P. W. 2019. Low rates of acquisition of de novo mutations in human pluripotent stem cells under different culture conditions.

Wilhelm, T., Olziersky, A. M., Harry, D., De Sousa, F., Vassal, H., Eskat, A. & Meraldi, P. 2019. Mild replication stress causes chromosome mis-segregation via premature centriole disengagement. Nat Commun, 10, 3585.

Zhang, J., Hirst, A. J., Duan, F., Qiu, H., Huang, R., Ji, Y., Bai, L., Zhang, F., Robinson, D., Jones, M., Li, L., Wang, P., Jiang, P., Andrews, P. W., Barbaric, I. & Na, J. 2019. Anti-apoptotic Mutations Desensitize Human Pluripotent Stem Cells to Mitotic Stress and Enable Aneuploid Cell Survival. Stem Cell Reports, 12, 557–571.

